# MicroRNA-222 regulates fluid shear stress-induced human nucleus pulposus cells degeneration through affecting c-Fos expression

**DOI:** 10.1101/524595

**Authors:** Haixiong Miao, Yicun Yao, Baoqing Ye, Libing Dai, Weiguo Liang

**Affiliations:** Guangzhou City Red Cross Hospital, Guangzhou Institute of Traumatic Surgery, The Fourth Affiliated Hospital of Medical College, Jinan University, Guangzhou 510220, China

**Keywords:** fluid shear stress, c-Fos, microRNA-222, nucleus pulposus cells, ERK5

## Abstract

Intervertebral disc degeneration (IVDD) is a chronic disease that correlates with the deterioration of the nucleus pulposus (NP) cells. However, the molecular mechanism of IVDD remains unclear. In this study, we investigated the function of microRNA-222 in IVDD and the potential molecular mechanism. NP cells treated with fluid shear stress (FSS) were used to simulate a model of IVDD in vitro. MicroRNA-222 was significantly downregulated in NP cells stimulated with FSS compared with that in unstimulated NP cells. Human NP cells were also treated with FSS to induce their degeneration. The mRNA and protein levels of C-FOS, MEK, phosphorylated MEK5 (pMEK5), ERK5, and pERK5 were evaluated with RT-PCR and western blotting, respectively. Enzyme-linked immunosorbent assays were used to investigate type II collagen and Aggrecan expression. NP cell proliferation was determined with the Cell Counting Kit-8. MicroRNA-222 was significantly downregulated in NP cells treated with FSS. The production of c-Fos and MEK5 were markedly reduced or increased in NP cells transfected with the has-microRNA-222 mimic or inhibitor, respectively, whether or not they were stimulated with FSS. The overexpression or inhibition of microRNA-222 markedly accelerated or suppressed the apoptosis of FSS-stimulated NP cells, respectively. In the NP cells, the overexpression or inhibition of microRNA-222 massively inhibited or strengthened Aggrecan and type II collagen expression. Together, our data indicated that c-Fos was a target of microRNA-222, and was negatively regulated by microRNA-222 in NP cells. Our findings also suggested that microRNA-222 is a possible therapeutic target for IVDD because it regulates c-Fos.

## Introduction

The intervertebral discs play a vital role in moving, bending, and supporting the weight of the spine. Intervertebral disc degeneration (IVDD) is one of the major health problems associated with low back pain and is a burden on societal economies[1]. Previous studies have demonstrated that the intervertebral disc has an asymmetric and complex organization, and that its metabolic activity is controlled by genetic factors, nutritional transmission disorders, environmental factors, and fluid shear stress (FSS) [2]. Several studies have suggested that nucleus pulposus (NP) cells maintain the integrity of the intervertebral discs by generating aggrecan, type II collagen, and other factors [3]. The stimulation of NP cells with FSS causes enormous changes in their water and proteoglycan contents [4]. Previous research has confirmed that FSS affects the disc metabolism by mechanically irritating the bone cells, which translate this irritation into cellular biochemical signals [5]. However, the exact molecular mechanism underlying the regulation of NP cell degeneration remains unclear. Understanding the mechanism that regulates the degeneration of NP cells will be important in treating IVDD in the future.

The early transcriptional processes after the extracellular stimulation of cells is controlled by the heterodimeric transcription factor activating protein 1 (AP1), which is composed of c-Fos, FOSB, or C-JUN[6]. Several research groups have reported that the activity of c-Fos correlates with MAPK pathways in various cell types [7]. However, the molecular mechanism that underlies the regulation c-Fos expression under FSS is still not clear.

MicroRNAs (miRNAs) are a novel class of functional noncoding RNAs that negatively control mRNA stability by combining with the 3 untranslated regions of their target mRNAs to facilitate mRNA degradation and/or block RNA translation[8]. In recent years, increasing studies have suggested that miRNAs regulate almost all cellular processes and are often aberrantly expressed in various physiological abnormalities and diseases. There is strong evidence that the abnormal expression of miRNAs, such as miRNA-210, miRNA-96, miRNA-494, and miRNA-223, is a common and crucial factor regulating NP cell degeneration in IVDD.[9–14] The miRNA-222 promotes the apoptosis of dendritic cells, thus playing an important role in thyroid papillary cancer, breast cancer, pancreatic cancer, hepatocellular carcinoma, and lung cancer [15–17]. To our knowledge, the function and mechanism of miRNA-222 in regulating the NP degeneration induced by FSS remains unknown. To investigate the molecular mechanisms affecting NP cells stimulated with FSS, we examined the functions of miRNA-222 in the activation of c-Fos and the degeneration of NP cells. Our results demonstrated the novel finding that the inhibition of miRNA-222 upregulated the level of c-Fos and activated the MEK–ERK5 pathway in NP cells, accelerating their degeneration.

## Materials and methods

### Ethics statement

The intervertebral disc samples, obtained during orthopedic surgery for scoliosis, were from seven patients (aged 6–15 years) treated at the Zhongshan Hospital Affiliated with the First Military Medical University. All the patients gave their informed consent to the medical experiments. The use of detached human intervertebral discs obtained during surgery complied with the relevant guidelines of the Ethics Committee of Zhongshan Hospital Affiliated with the First Military Medical University.

### Intervertebral disc harvesting and NP cell isolation

The intervertebral disc samples were collected with sterile techniques. The gelatinous NP tissue was identified and dissected from the inner part of the intervertebral disc, mechanically minced, and digested in 0.25% type II collagenase (Sigma-Aldrich, USA) for 16 h at 37°C. The suspended cells were filtered through a 200-mesh filter and then centrifuged, washed, and seeded in 25 cm2 cell culture flasks at 3 × 105 cells/mL. The cells were stored in DMEM-F12 (Invitrogen Life Technologies, Carlsbad, CA, USA) containing 10% fetal bovine serum (Invitrogen Life Technologies), 0.5% gentamicin (Life Technologies), and 1% penicillin and streptomycin (Life Technologies), and incubated under 5% CO2 at 37°C. The culture medium was replaced every other day.

### Cell loading

Frangos et al. was built the fluid flow. Parallel glass plates coated with polydimethylsiloxane (PDMS) were used to make microfluidic flow chambers [18]. The shear stress on the laminar wall was calculated with the Poiseuille law: t = 6Qμ/wH2, where Q is the flow rate (0.833 cm3/s), μ is the viscosity of the medium (0.003 dyn s/cm2), w is the width of the flow chamber (2.0 cm), and h is its height (2.0 cm) [19, 20]. During the FSS experiment, the collected NP cells were cultured in 6-well plates pretreated with 10 μg/mL fibronectin at 37°C under 5% CO2 until they were approximately 80% confluent. The flasks were exposed to 12 dyn/cm2 fluid shear for 45 min, according to previous research. Unloaded controls were cultured under equivalent conditions without fluid flow.

### Cell transfection

NP cells were seeded in 24-well plates and cultured to 40%–50% confluence. The cells were then transfected with the has-mimic222 (mimic222), has-inhibitor222 (inhibitor222), or has-negative control (NC) (control) (GenePharma, Shanghai, China) using Lipofectamine 3000 (Invitrogen, Carlsbad, CA, USA), according to the manufacturer’s instructions. The final concentration of mimic used was 50 nM. Untransfected cells and cells transfected with a scrambled RNA sequence were used as controls. The sequences used were as follows:has-microRNA-222 mimics sense,5’-UUCUCCCAACGUGUCACGUTT-3’;antisense,5’-CCAGUAGCCAGAUG UAGCUUU-3’; has-microRNA-222 inhibitor, 5’-ACCCAGUAGCCAGAUGUAGCU-3’.

### Reverse transcription-quantitative real-time PCR (RT-PCR) analysis

The total RNA of NP cells was or was not treated with FSS, and then transfected with the mimic222 or inhibitor222, and was isolated with TRIzol reagent (Invitrogen). The First Strand cDNA Synthesis Kit (Takara) was then used to reverse transcribe it into cDNA. The qPCR was performed on a 7500HT Fast Real-Time PCR System (Applied Biosystems, Singapore). The sequences of the primers were as follows:U6 sense 5’-CGCTTCGGCAGCACATATAC-3’,antisense 5’-CAGGGGCCATGCTAATCTT-3’:microRNA-222 sense 5’-GCGAGCTACATCTGGCTACTG-3’, antisense 5’-GTGCAGGGTCCGAGGT-3’;c-Fos sense 5’-GTGCCCTATGGCGAATTCAA-3’,antisense 5’-GCACGTGTTCCAGTGTGAGG-3’

### Western blotting analysis

After each treatment, the NP cells were washed twice with ice-cold phosphate-buffered saline and then lysed in RIPA lysis solution (50 mM Tris [pH 7.4], 1% Nonidet P-40, 0.1% SDS, 0.5% sodium deoxycholate, 150 mM NaCl, and protease inhibitors). A BCA kit (Beyotime, China) was used to determine the concentrations of the protein. Equal amount of protein were detected with SDS-PAGE and transferred to nitrocellulose membranes (Bio-Rad Laboratories), which were then incubated with the primary antibodies ERK5(#3372,CST,1:1000),Phospho-ERK5(Thr218/Tyr220)(#3371,CST,1:1000),c-Fo s(#4384,CST,1:1000),MEK5(ab45146,Abcam,1:5000),phospho-MEK5(S311/T315)(# PA5-37701,Thermo Fisher,1:1000), at 4°C overnight. After three washes in Tris-buffered saline containing Tween 20 (TBST), the membranes were incubated at 37°C for 2 h with the corresponding secondary antibody: horseradish peroxidase (HRP)-linked anti-mouse IgG antibody (#7076, Cell Signaling Technology [CST]; diluted 1:2000) or HRP-linked anti-rabbit IgG antibody (#7074, CST; diluted 1:2000). An anti-β-actin antibody (#4967, CST; diluted 1:500) was used as the internal loading control. The ImageJ software (GE, Piscataway, NJ) was used to measure the gray values.

### Cell proliferation assay

Briefly, cell proliferation was analyzed with Cell Counting Kit-8 (CCK-8, Beyotime, China). The differently treated NP cells were seeded in 96-well plates at 3 × 103 cells/well, cultured for 0, 24, 48, or 72 h. Each well was treated with CCK-8 solution and the absorbance read at 450 nm. The data are presented as the percentage cell viability in the differently treated groups. All experiments were repeated in triplicate.

### Enzyme-linked immunosorbent assays (ELISAs)

ELISA kits (USCN Life Science, Wuhan, China) were used to detect the expression of Type II collagen and Aggrecan in the extracellular fluid, according to the manufacturer’s instructions.

### Statistical analysis

All results were tested in three independent cultures for each experiment. Data were presented as the means±SD. SPSS 20.0 software was analyzed the data. The statistical significance was calculated by Student’s t-test or one-way analysis of variance followed by Duncan’s multiple comparison test. p < 0.05 was considered as significant

### Authorship

HXM and YCY were designed the study. LBD was acquisitioned the data. BQY was analyzed the data. HXM was drafted the article. WGL was final approval of the version to be submitted.

## Acknowledgments

This work was supported by Guangzhou Scientific and technological project of combining traditional Chinese medicine with Western medicine (20182A010012); Guangdong Medical Scientific Research Foundation (A2016520, A2017501); Guangdong Natural Science Foundation (2015A030313736) Guangdong Administration of Traditional Chinese Medicine (20171207);Guangdong Science and Technology Planning Project (2013B021800070, 2012B031800338); Guangzhou Medial Science and Technology project (20171A011252, 20161A011017, 20161A011016). The authors have no conflicts of interest.

## Results

Compared with the control NP cells, 89 miRNAs were upregulated and 97 miRNAs were downregulated in the NP cells treated with FSS. Because the signal at both sides being greater than eight and the multiple between them being greater than two, miRNA-222 was chosen for further study with a bioinformatics analysis and a critical review of the literature. The predicted target gene of miRNA-222 was a member of the ERK family.

### Upregulation of miRNA-222 and extracellular signaling-regulated kinase 5 (ERK5) signaling activity in FSS-stimulated NP cells

MiRNA-222 is expressed in diverse tissues, and anomalous miRNA-222 expression has been associated with several diseases. To determine the biological roles of miRNA-222 in NP cell degeneration, a model of NP cell degeneration was generated with FSS in this study. We compared the expression of miRNA-222 in human NP cells treated with FSS or not using RT-PCR (Fig. 1A). The miRNA-222 expression was massively lower in the NP cells transfected with a scrambled microRNA(control) treated with FSS than in the cells not treated with FSS, suggesting that miRNA-222 played a vital role in NP cell degeneration. Interestingly, western blotting indicated that MEK5 was reduced (Fig. 1B), whereas pMEK5, pERK5, and c-Fos were increased (Fig. 1C–E), and ERK5 was unchanged (Fig. 1F) in the FSS-treated cells. These data suggested that miRNA-222 upregulated the MEK5 pathway in FSS-stimulated NP cells. However, the mechanism was unclear.

**Figure 1.**
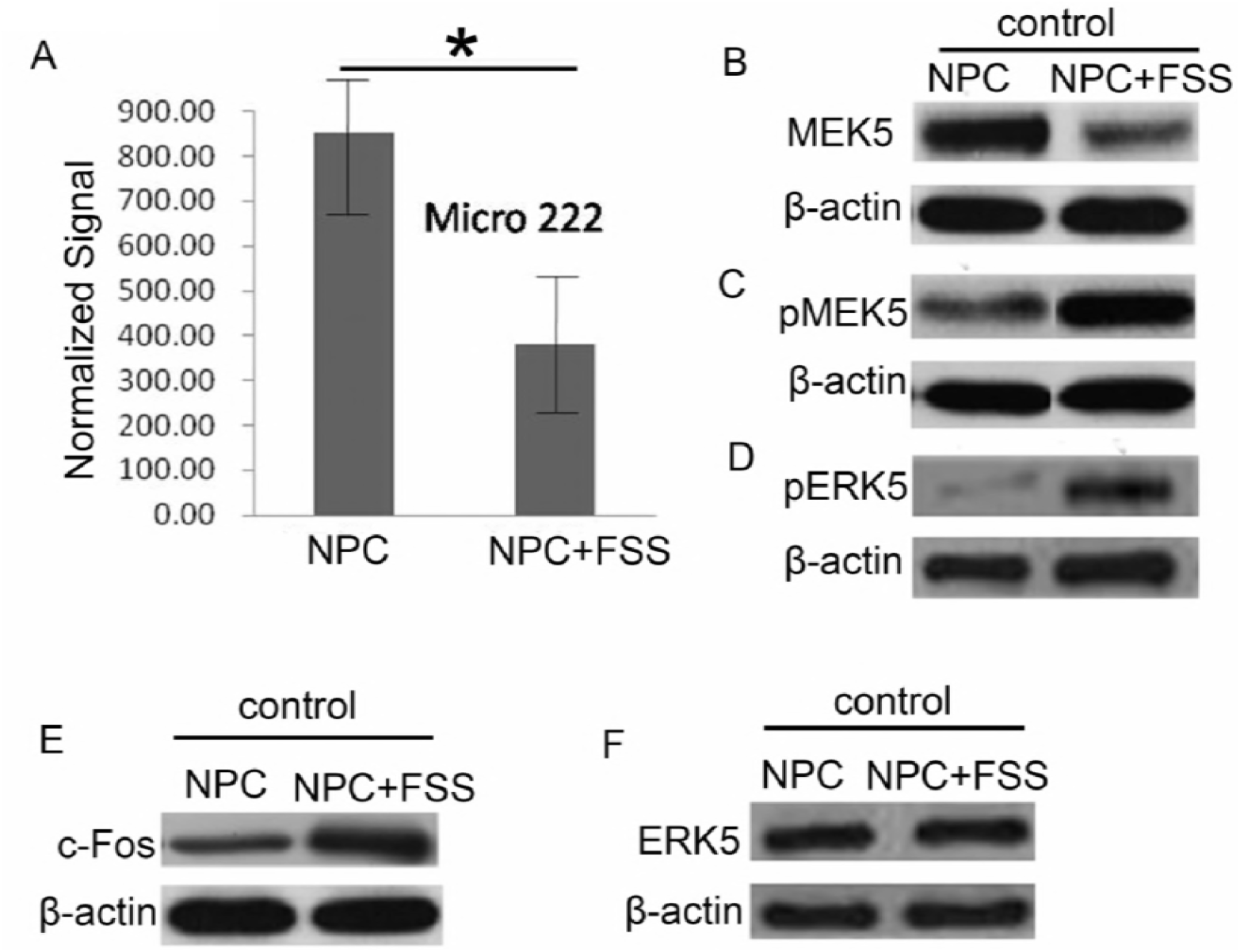
Upregulation of miRNA-222 and ERK5 signaling activity in FSS-stimulated NP cells. (A) RT-PCR analysis of miRNA-222 expression in FSS-stimulated NP cells. (B–F) Protein levels of MEK5, pMEK5, pERK5, c-Fos, and ERK5 in control group with FSS-stimulated or not, determined with western blotting. (*n* = 6). \**p* < 0.05

### MiRNA-222 inhibits the activity of the MEK–ERK pathway in FSS-stimulated NP cells

To investigate the regulatory effect of miRNA-222 on IVDD, NP cells were activated or blocked by transfecting them with a mimic222 or inhibitor222, respectively. NP cells transfected with a scrambled microRNA were used as the control. The cells were then stimulated with FSS. At 45 min after stimulation, miRNA-222 expression was analyzed with RT-PCR (Fig. 2A–B), and the levels of MEK/ERK-related proteins were detected with western blotting. Compared to the control group, the mimic222 significantly reduced the levels of pMEK5 and pERK5 (Fig. 2C–D). Interestingly, MEK5 expression was also dramatically reduced (Fig. 2E), whereas the protein level of ERK5 remained unchanged (Fig. 2F). Conversely, the inhibitor222 significantly increased the levels of pMEK5 and pERK5 (Fig. 2G–H), and dramatically increased the expression of MEK5 (Fig. 2I). However, the protein level of ERK5 remained unchanged (Fig.2J). These data indicated that the downregulation of miRNA-222 significantly reduced MEK/ERK expression in FSS-stimulated NP cells.

**Figure 2.**
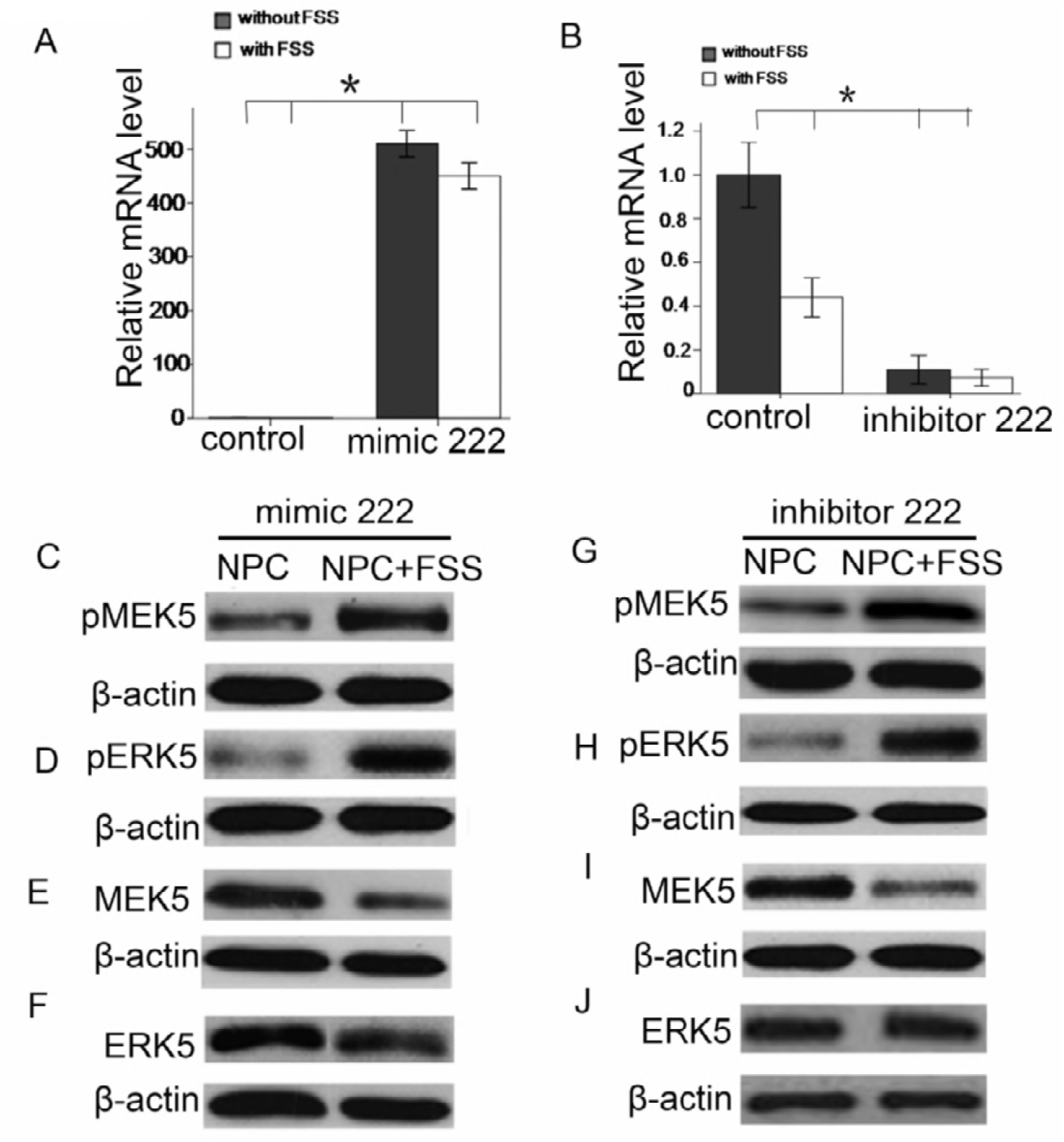
MiRNA-222 inhibits the activity of the MEK–ERK pathway in FSS-stimulated NP cells. (A) RT-PCR analysis of miRNA-222 expression in NP cells transfected a mimic222 and treated with or without FSS. (B) RT-PCR analysis of miRNA-222 expression in NP cells transfected with an inhibitor222 and treated with or without FSS. (C–F) Western blotting analysis of pMEK5, pERK5, MEK5, and ERK5 levels in NP cells transfected with a mimic222 and treated with FSS or not. (G–J) Western blotting analysis of pMEK5, pERK5, MEK5, and ERK5 levels in NP cells transfected with an inhibitor222 and treated with FSS or not. (*n* = 6). \**p* < 0.05

### MiRNA-222 inhibits the activity of c-Fos and affects the MEK–ERK-c-Fos pathway in NP cells

A predictive analysis indicated that c-Fos was a target gene of miRNA-222. It has been reported that c-Fos is involved in cellular immunity and the inflammatory response [15,16]. Our results suggested that FSS did not affect the expression of ERK5 in NP cells. Therefore, we speculated that other factors affected the MEK-ERK5-C-FOS signaling pathway in IVDD, so we investigated whether miRNA-222 regulated c-Fos expression in NP cells. Following the transfection of NP cells with a mimic222 or inhibitor222 for 24 h, we then treated the cells with or without FSS (12 dyn/cm2 × 45 min), and detected the levels of c-Fos protein with western blotting analysis. Compared to the control group, c-Fos expression was massively reduced after the upregulation of miRNA-222 (Fig. 3A), whereas the levels of MEK5, pMEK5, ERK5, and pERK5 were unaffected (Fig. 3B–E). Conversely, when miRNA-222 expression was downregulated by the inhibitor, the protein levels of c-Fos increased massively (Fig. 3F), whereas the levels MEK5, pMEK5, ERK5, and pERK5 remained unchanged, compared with the control group which cells stimulated with FSS or not(Fig. 3B–E). Interestingly, protein quantification also indicated that MEK5 expression was dramatically increased in the mimic222 group. However, the mRNA levels of c-Fos did not differ across the control group, the mimic-treated group, and the inhibitor-treated group, with or without FSS (Fig. 3G). And c-Fos protein expression was dramatically reduced in the mimic222 group (Fig. 3G). These data indicated that miRNA-222, not FSS, regulated the expression of c-Fos and affected the MEK-ERK-c-Fos pathway. Generally, these results showed that the inhibition miRNA-222 significantly induced the expression of c-Fos in FSS-stimulated NP cells and activated the MEK-ERK-c-Fos pathway. These findings indicated that miRNA-222 exerted an inhibitory effect on c-Fos expression after its transcription, indicating that c-Fos was a newly identified target protein of miRNA-222 and that miRNA-222 might offer a new therapeutic target for IVDD.

**Figure 3.**
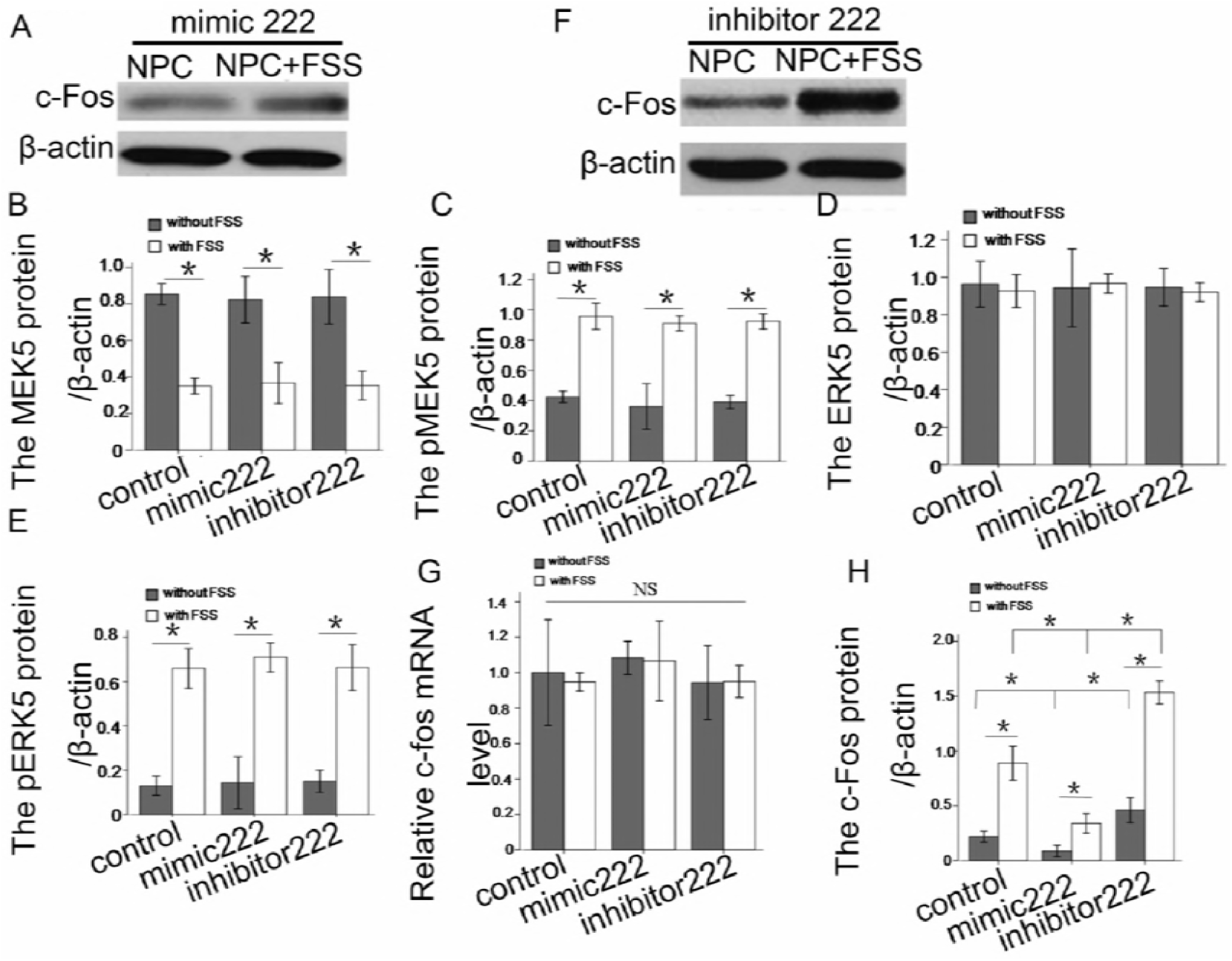
MiRNA-222 inhibits the activity of c-Fos and affects the MEK-REK-c-Fos pathway in NP cells. (A) Western blotting analysis of c-Fos expression in NP cells transfected with a mimic222 and treated with FSS or not. (B–E) Protein quantification of MEK5, pMEK5, ERK5, and pERK5 in the control group, mimic222 group, and inhibitor222 group, treated with FSS or not. (F) Western blotting analysis of c-Fos expression in NP cells transfected with an inhibitor222, and treated with FSS or not. (G) RT-PCR detection of c-Fos mRNA levels in the control group, mimic-transfected group, and inhibitor-transfected group, treated with or without FSS. (H) Quantification of c-Fos protein in the control group, mimic-transfected group, and inhibitor-transfected group, treated with or without FSS. (*n* = 6). \**p* < 0.05, \*\**p* < 0.01

### MiRNA-222 accelerates NP cell degeneration

A CCK8 assay was used to determine the proliferation of NP cells transfected with a mimic222 or inhibitor222 to examine the underlying function of miRNA-222. Compared with the control group, cell proliferation was reduced by the transfection of the mimic222 and increased by the transfection of the inhibitor222 (Fig. 4A). These data suggested that miRNA-222 played a significant role in controlling NP cell proliferation.

**Figure 4.**
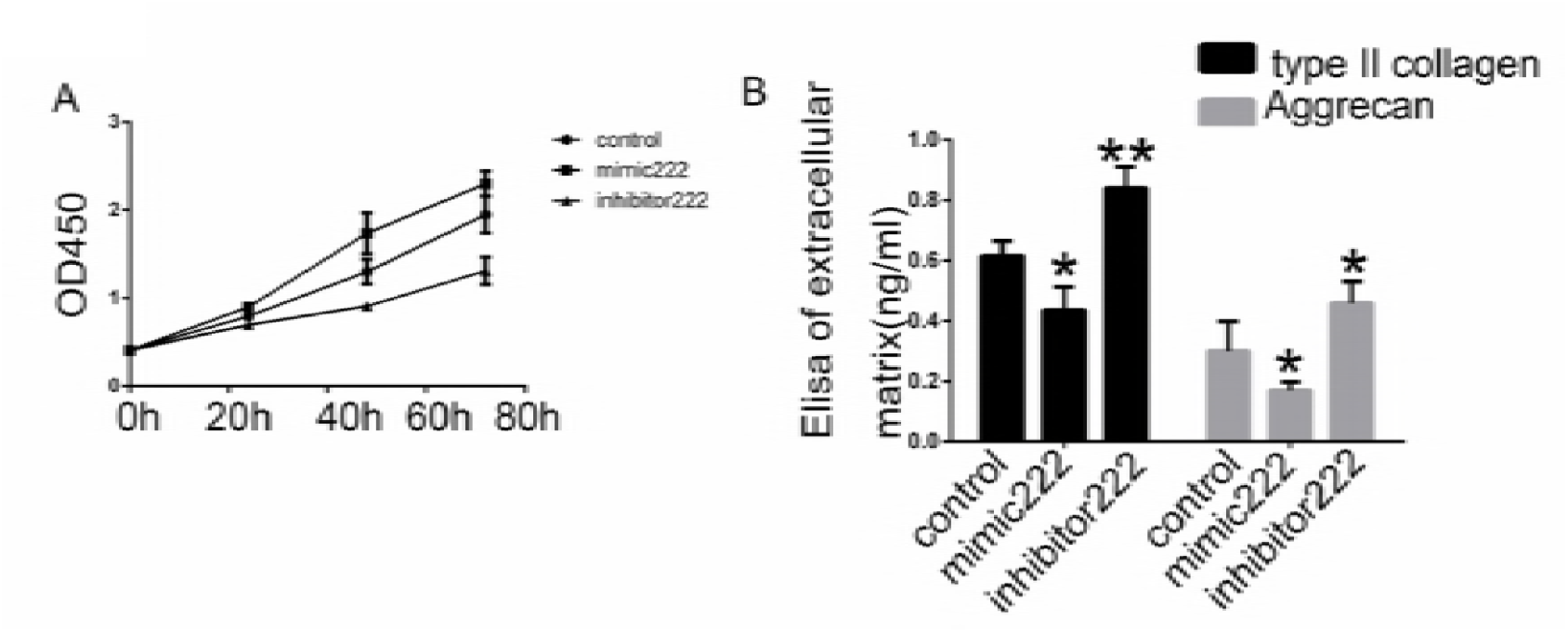
MiRNA-222 accelerates NP cell degeneration. (A) CCK8 assay of cell proliferation after NP cells were transfected with the mimic222 or inhibitor222, and treated with or without FSS. (B) ELISA analysis of type II collagen in NP cells transfected with control, mimic222 or inhibitor222.

ELISAs confirmed the reduction of type II collagen and Aggrecan expression in the FSS-treated NP cells relative to that in the control NP cells. The miRNA-222 mimic clearly suppressed the expression of type II collagen and aggrecan, whereas the miRNA-222 inhibitor significantly increased the levels of type II collagen and aggrecan compared with their levels in cells only treated with FSS (Fig. 4B).

Thus, we have demonstrated that NP cells degenerated when the MEK-ERK pathway was activated. The inhibition of miRNA-222 reduced the expression of collagen II and Aggrecan and induced c-Fos expression. Therefore, miRNA-222 may have affected NP cell degeneration by targeting the c-Fos gene.

## Discussion

In this study, we established a model of NP cell degeneration in the physiological state, in which external mechanical stress (FSS) was applied to human NP cells. MiRNA-222 expression levels and functions were investigated in these FSS-treated NP cells. We confirmed that miRNA-222 expression was inhibited in NP cells treated with FSS. The abnormal ectopic expression of miRNA-222 accelerated NP cell degeneration. Conversely, the inhibition of miRNA-222 induced MEK5 and ERK5 activity, and increased the levels of pMEK5, pERK5, and C-FOS in NP cells treated with FSS. In this study, c-Fos was shown to be an immediate target of miRNA-222 in NP cells. The overexpression miRNA-222 reduced the levels of c-Fos, MEK5, and pERK5 and enhanced NP cell proliferation. Similarly, the ectopic expression of miRNA-222 increased the levels of MEK5 and pERK5 by activating c-Fos in NP cells. These data suggested that miRNA-222 affected the evolution of IVDD.

The cytoskeleton prevents the mechanical load response inside cells [21]. The activity of NP cells is controlled by the mechanical loading on the intervertebral discs. Our research has shown that miRNA-222 promoted the proliferation of NP cells by targeting c-Fos and activating its downstream signaling molecule ERK5, which is a member of the mitogen-activated protein kinase (MAPK) family. MEK5 is the only upstream kinase to activate the expression of ERK5. Previous research has shown that ERK5 is involved in the cellular responses to growth factors and hypoxia inducible factor 1 (HIF-1), cytoskeletal organization, proliferation, and chemotaxis [22, 23]. The autophosphorylation of the C-terminal domain of ERK5 indicates that it differs from other members of the MAPK family in exerting its biological functions and controlling the related mechanisms [24].

c-Fos was identified as an immediate target gene of miRNA-222 in this study. Research into the proto-oncogene c-Fos and its product, the c-Fos protein, has recently become very important. c-Fos is used extensively as a marker of cellular responses to stimuli. Interesting research conducted by Kostenuik showed that after rat hind limbs were elevated for 5 days and the bone-marrow stromal cells (BMSCs) from the hind limbs were harvested and cultured, the c-Fos expression in these cells was significantly reduced [26]. The expression of c-Fos was also reduced in cultured BMSCs from rat hind limbs that had been elevated for 5 days. Previous studies have shown that the function of skeletal unloading on bone formation in the growing rat is transient. Therefore, the level of c-Fos controls bone formation when the skeleton is unloaded. It can be speculated that c-Fos acts as the on-off switch for a nuclear trigeminal messenger and a gene transcriptional regulator.

The miRNAs are involved in the diagnosis and prognosis of numerous diseases and are latent targets for gene therapy. MiRNA-222, located on chromosome Xp11.3, is a member of the miRNA-221/222 family and always acts as a gene cluster in cellular regulation[27, 28]. It has been suggested that miRNA-222 could be used as a therapeutic tool to regulate cell proliferation or modulate cell sensitivity to anticancer agents. Studies have shown that miRNA-222 is downregulated during the normal growth and development of red blood cells. MiRNA-222 is also associated with colorectal cancer. Another study indicated that miRNA-222 directly targets c-Fos to regulate the levels of protein expression in cutaneous melanoma [29]. However, how miRNA-222 is expressed and functions in IVDD are not fully understood. Interestingly, miRNA-222 expression was reduced in NP cells treated with FSS in this study. We also showed that the inhibition of miRNA-222 activated c-Fos activity and accelerated NP cell degeneration.

In conclusion, miRNA-222 promoted the proliferation of human NP cells by regulating c-Fos expression, increasing the phosphorylation of the c-Fos protein, and activating the MEK-ERK pathway, suggesting that miRNA-222 was a crucial factor in the pathogenesis of IVDD. Our data highlighted the importance of studies of the roles of miRNA in regulating the NP cell degeneration induced by FSS. This study confirmed that the downregulation of miRNA-222 enhanced c-Fos expression. On the contrary, the overexpression of miRNA-222 inhibited the expression of c-Fos. These findings suggested that mRNA-222 is a new therapeutic target for the diagnosis and treatment of NP cell-related diseases.

